# DNA from *cetariae* fish remains confirms sardine (*Sardina pilchardus*) use and local population continuity in Northwestern Iberia since Roman times

**DOI:** 10.1101/2025.01.28.635260

**Authors:** Gonçalo Espregueira Themudo, Adolfo Fernández-Fernández, Patricia Valle Abad, Alba A. Rodríguez Nóvoa, Carlos Fernández-Rodriguez, Eduardo González-Gómez de Agüero, Rute R. da Fonseca, Paula F. Campos

## Abstract

The Romans were among the first to extensively exploit fish resources, establishing large-scale salting and preservation plants. Small pelagic fish were fermented to produce sauces like garum. Here, we recover DNA from specimens collected at Adro Vello (O Grove), Galicia, dating to the 3rd century AD. Whole genome resequencing, confirmed these as European sardines (*Sardina pilchardus*) and assigns all samples to a population ranging from North Morocco to the Bay of Biscay. The low admixture levels in ancient samples suggest lower connectivity in Roman times. This overlooked archaeological material can enhance our understanding of past subsistence economy, culture, and diet.

## Introduction

Marine resources have been vital for humans, providing an alternative food, fuel, clothing and various raw materials, significantly shaping the economic trajectories of European societies. Fish was a vital source of protein in ancient Rome, and given the scarcity of refrigeration, mainly used for the commerce of oysters, the preservation of fish was crucial both for local consumption and trade. Therefore, Romans established fish salting plants, also known as “cetariae” in coastal areas throughout the Roman Empire. These facilities processed and preserved fish, mainly through salting and fermentation (Bernal-Casasola *et al*. 2018). Large fish, such as tuna, were cleaned, gutted, and layered with salt in stone vats. This salting process drew out moisture, preventing bacterial growth allowing the fish to be stored for long periods and facilitated its’ commerce throughout the Roman Empire.

Small fish were not considered of great quality and therefore were used in the making of fish sauces, of which the most famous is *garum*, used for seasoning and as condiments, conferring an umami flavour to food. *Garum*, *liquamen*, *allec* or *muria* (Grainger 2014) were prepared by crushing the complete fish and then fermenting it in brine. So, before the relatively well documented exploitation of the past decades, pelagic fish, like sardines, sprats, anchovies and mackerels were already a very important component of human diet in the form of fish sauce. Sardines were also used for the making of fish pastes, that beyond fish could contain other marine fauna and sometimes even meat (Marzano 2018). The trade of fish sauce from Hispania and Lusitania into the rest of the Roman Empire, in characteristic amphorae, was a flourishing activity and is extremely well documented (Fernández 2014).

Fish bones are very common in the archaeological record, however their use in archaeogenomic studies has been minimal, probably due to its fragmentary nature which makes it difficult to identify species or even genus. Ancient DNA from fish remains has however been recovered from a multitude of different species (Oosting *et al*. 2019). Their potential is enormous, as their assemblies open a new research avenue to understanding subsistence economy, culture, and the diet of past peoples.

The use of ancient DNA (aDNA) techniques allows us to explore the past and is able to provide us with a range of information that cannot be obtained using fishery catch data or modern specimens alone, like demographic dynamics and adaptive change over time. Studies have not only focused on species with a considerable size like catfish (Arndt *et al*. 2003), bluefin tuna (Andrews *et al*. 2021) or cod (Star *et al*. 2017; Pinsky *et al*. 2021; Martínez-García *et al*. 2021) but also herring (Atmore *et al*. 2022, 2024) and have shown good preservation and high levels of endogenous DNA in most remains (Star *et al*. 2017; Oosting *et al*. 2019; Ferrari *et al*. 2021) even with small amounts of starting material (Atmore *et al*. 2023).

The bottom of fish salting vats offers a myriad of remains from where DNA could be retrieved. One of the biggest challenges of looking into pelagic fish from the bottom of fish salting vats is their small size and the small size of the bone material available. Due to the crushing of the specimens to make the fish sauce/paste, most bones are non-articulated, and broken vertebrae are the most common bone present. Chances of retrieving complete specimens in these archaeological sites are thus hampered by such practices, and fragile and fractured bones are often difficult to identify to species level. The small bone size also limits the initial amount of tissue for DNA extraction and library construction.

DNA preservation in these samples may be affected by the fermentation process and other food processing they went through. Fermentation leads to lower pH, and increase in the depurination rate and hence fragmentation of the DNA molecule (Bauer et al. 2003). The proliferation of microorganisms in this environment, and its’ endonucleases, will also lead to higher levels of DNA degradation (Gryson 2010; Garnier *et al*. 2018). Processes such as grinding and maceration accelerate DNA degradation (Gryson 2010).

Moreover, specimens in the brine were in contact with other sardines or even other pelagic fish, potentially contaminating samples with DNA from the same species, i.e., endogenous DNA from the sardine bone and contaminant DNA from the sardines in the brine.

Here we attempt to extract DNA, build double strand DNA libraries and sequence small bone remains (mainly crushed vertebrae), morphologically identified as European sardine (*Sardina pilchardus*) from the bottom of a Roman *cetariae* (workshop) in Adro Vello, O Grove, Galicia, where fish sauces and pastes were produced between the 1st century and the end of 3rd century AD, in order to access their suitability for aDNA research.

Adro Vello is an important archaeological site on the shore of O Carreiro beach. It has a long history of occupation from the Roman era to the 18^th^ century (Figure 1C). In chronological order we can find remains of the fish-salting plant (Figure 1B), a Roman villa, a possible early medieval monastery with a church and associated necropolis from the 7th to the 11th century, which was replaced by a medieval church and a large burial cemetery from the 13th century until the 18th century when the church was moved inland (Mangas-Carrasco et al., 2022). Within the Roman assemblage, four vats used for making fish sauces and pastes were found and in the bottom of vat number one a myriad of ichthyological fish residues were preserved. Some of these were sampled to assess the DNA preservation of such remains and its potential use in species identification.

**Figure 1.**
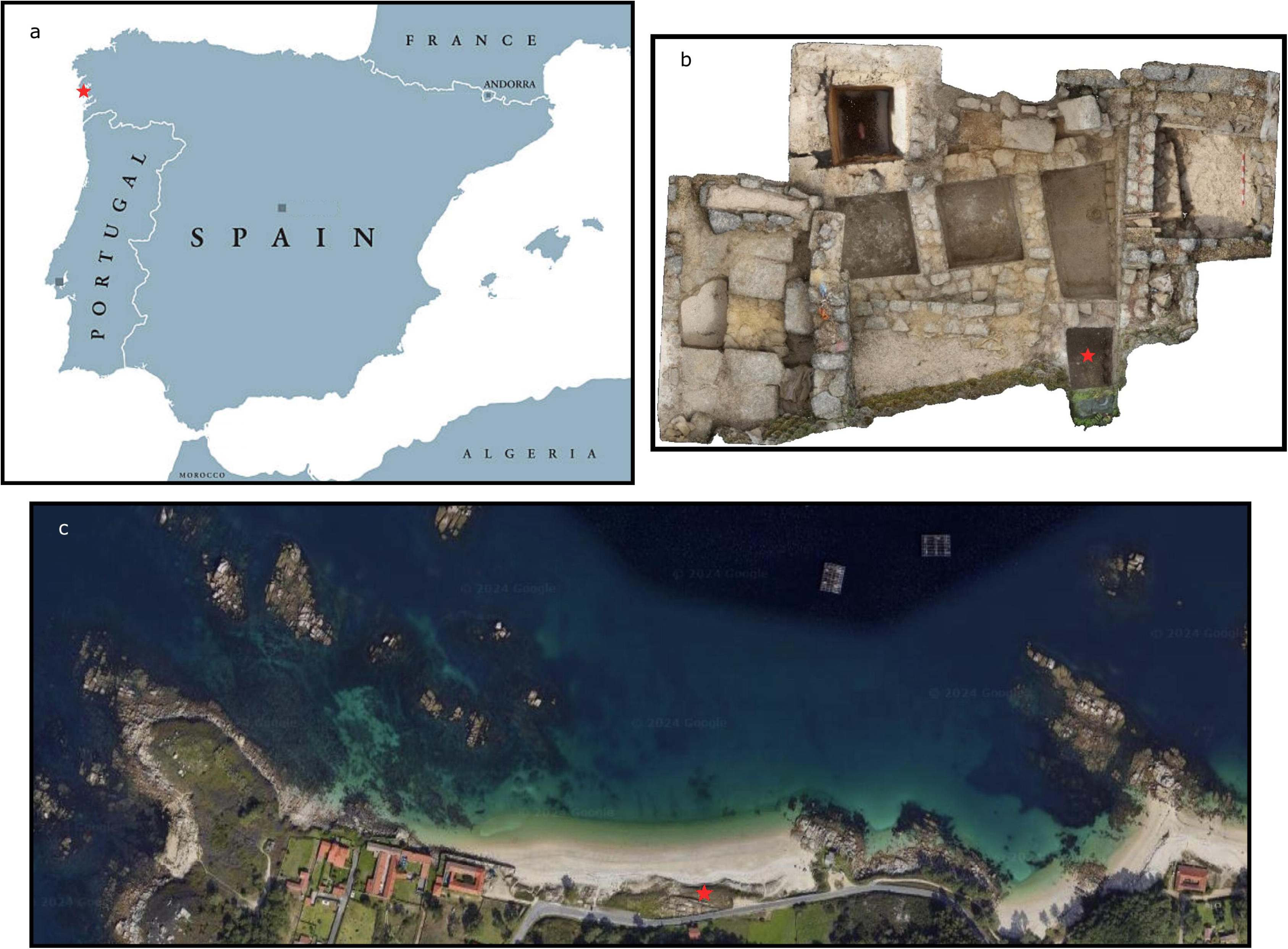
A Map of the Iberian Peninsula with the geographical location of Adro Vello, O Grove in red; B-3D reconstruction of the fish salting plant at Adro Vello, vat 1 marked with a red star; C – Aerial view of O Carreiro beach, the archaeological site of Adro Vello is marked with a red star.

This study aimed to determine whether shotgun sequencing of DNA extracted from small bones retrieved from *garum* could identify the fish species used in the production site and to assess the potential for DNA preservation in fish bones from fermentation brines.

Despite the clear potential of these remains, to our knowledge, no genomic studies have so far taken advantage of them to study past fish consumption and population dynamics of commercially relevant species.

## Material and Methods

### (a) Sampling, stratigraphic and radiocarbon dating

Ichthyological remains (Figure 2) were collected from the bottom of fish salting vat number one in the Roman site of Adro Vello, O Grove, Galicia, North-western Iberia (Figure 1). They were placed in zip lock bags, stored and processed in Laboratorio de Prehistoria, University of León. They were firstly washed through a sieve (0,8mm), separated into groups and anatomically and morphologically screened and identified as *Sardina pilchardus* (González Gómez de Agüero 2014).

**Figure 2.**
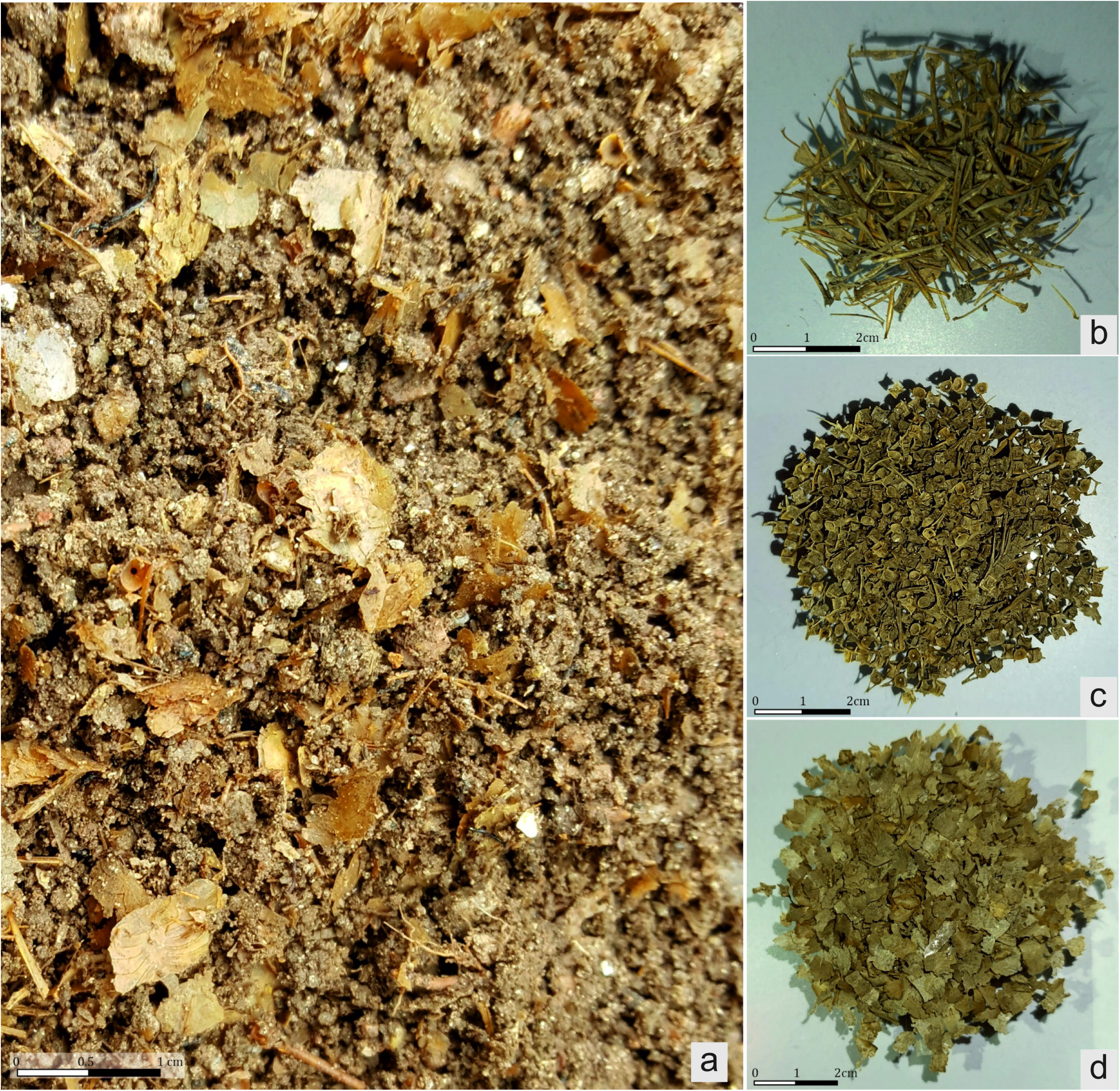
– A-Ichthyofauna remains from the bottom of vat1 before processing; B-fish spines; C-Fish vertebrae; D-Fish scales.

**Figure 3.**
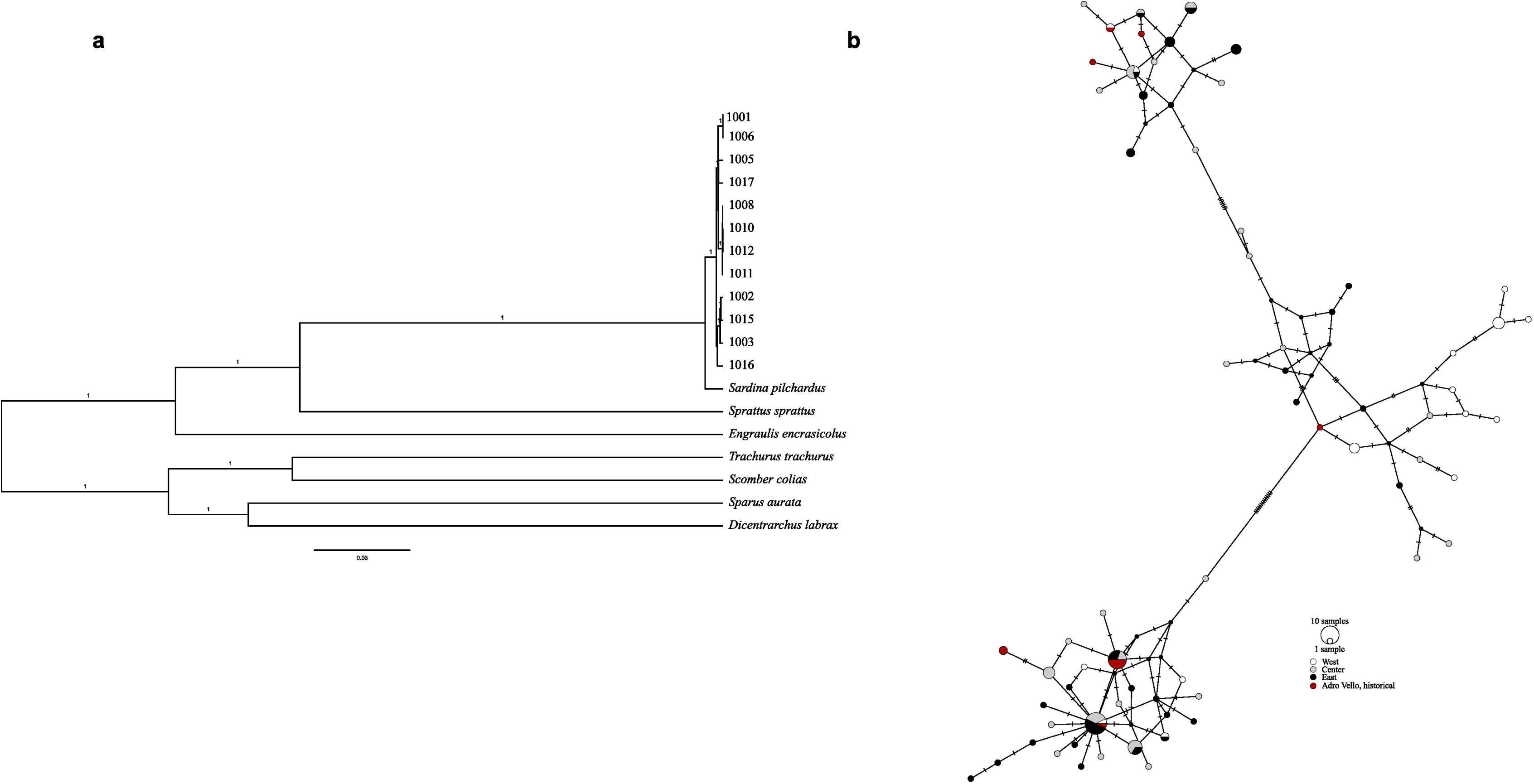
A-Bayesian phylogenetic reconstruction based on the complete mitochondrial genome of 12 historical sardine samples. Sequences of European sardine (NC009592); the Atlantic horse mackerel (NC006818); the Atlantic chub mackerel (NC013724), sprat (NC009593); anchovies (NC009581); European seabass (NC026074) and Gilthead seabream (NC024236) were also included in the analyses. Posterior probabilities above 0,9 are shown. B – Median joining network (PopArt) of 108 modern and 12 historical mitogenomes. Only variants with minor allele frequency above 25% were used, Hatch marks represent mutations. Colours represent the main ancestry of each individual (as defined in da Fonseca et al 2024) except for the Adro Vello historical samples, depicted in burgundy.

**Figure 4.**
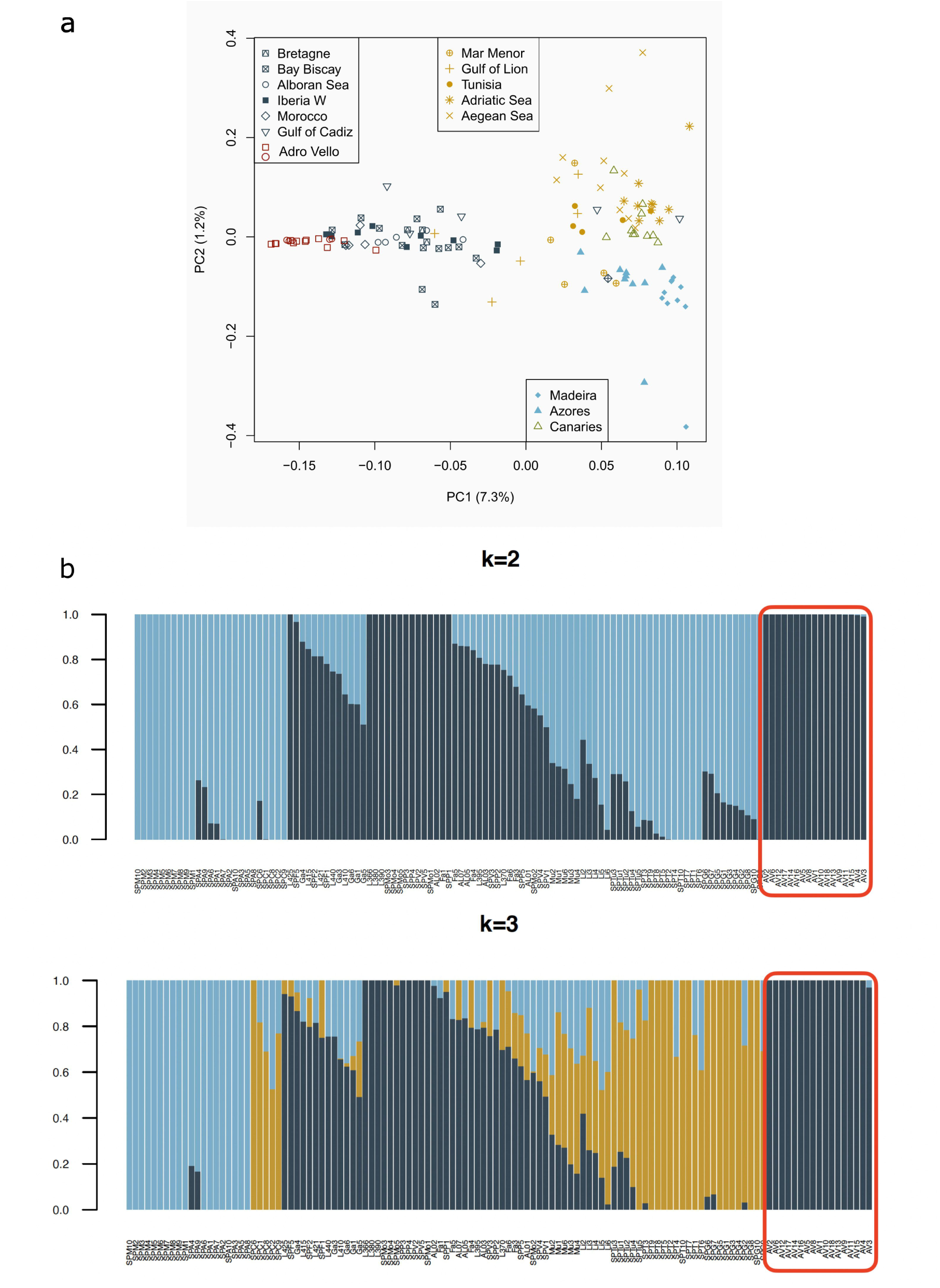
– A-Distribution of sardine populations based on the first three components of the Principal Component Analyses (PCA). Variation explained by each component is shown in parenthesis. Samples from the Western group are depicted in light blue, Centre group samples are in dark blue, Mediterranean samples in yellow. The samples from the Canary Islands are shown in green and the historical Adro Vello samples in burgundy (the five samples with lowest depth are depicted as circles). B-Population structure plot, obtained with NGSadmix, showing the ancestry of each individual (vertical bar) to two (above) and three (below) genetic clusters. Samples from Adro Vello are highlighted with a burgundy outline.

To confirm the age in which this salting plant was active a fish sample was sent to Beta Analytic for Accelerator mass spectrometry (AMS) radiocarbon dating (^14^C). Dates were calibrated with BetaCal4.20 and the High Probability Density Range Method (HPD): MARINE20 was used to correct for the local ocean reservoir effect (Heaton *et al*. 2020). Prior to analysis, bones were treated with alkalis in order to extract collagen.

### (b) DNA extraction and library building

All DNA extractions and double strand DNA Illumina library preparation steps were carried out in a dedicated ancient DNA facility at the Natural History Museum of Stockholm, University of Stockholm, Sweden, designed for dealing with potentially degraded samples such as these, which are particularly susceptible to contamination from exogenous sources of DNA. Contamination was monitored during the extraction and PCR processes by blank controls. Total cellular DNA from 18 bone samples, including vertebrae and opercula, was extracted from putative sardine specimens according to the following protocol: sardine bones, mainly vertebrae and opercula, were either crushed with a mortar or simply incubated overnight (smaller vertebrae) at 37 °C in 1.0 mL of 0.5 M EDTA and 25 mg/ml proteinase K. To pellet the non-digested powder, the solution was centrifuged at 12,000 rpm for 5 min. The liquid fraction was then transferred into a Centricon micro-concentrator (30-kDa cut-off), and spun at 4000 rpm for 10 min. When the liquid was concentrated down to 200–250 μl, the DNA was purified using the MinElute PCR purification kit (Qiagen), with the following modifications: a) 13× PB buffer (Qiagen) was used for the DNA binding step; b) spins were done at 8,000 rpm with the exception of the final one at 13,000 rpm; c) in the elution step, spin columns were incubated in 25 μl EB buffer at 37 °C for 10 min, spun down, and repeated once more. The eluates from both rounds of elution were pooled.

### (c) Library building and amplification

Double stranded DNA Illumina libraries were built from the extracts. A blunt-end library was constructed on 21.25 μl of DNA extract following the protocols outlined by Maricic et al. (2010) and Meyer and Kircher (2010). The amplified libraries were run on Agilent 2100 Bioanalyzer High Sensitivity DNA chip. The sequencing libraries were then pooled and screened at Novogene (150PE read mode). The 12 libraries with the highest % of endogenous DNA were then deep sequenced in one fourth of a lane on a llumina Novaseq X (150 cycles, PE read mode) at SciLife Lab, Sweden.

### (d) Data processing, mapping and annotation

Base calling was performed using the Illumina software CASAVA 1.8.2, with the requirement of a 100% match to the 6-nucleotide index used during library preparation. Raw Illumina reads were first processed with Trimmomatic (version 0.36) (Bolger *et al*. 2014) for removal of adapter sequences and trimming bases with quality < 20 and discard reads with length < 30. Clean reads were mapped to the European sardine genome assembly using bwa-mem version: 0.7.17-r1188 (Li 2013) and samtools version: 1.7 (Li *et al*. 2009) was used to retain reads with mapping quality >25. PCR duplicates were removed with Picard MarkDuplicates (version 1.95; http://picard.sourceforge.net).

The mitochondrial genome for each individual was obtained as a consensus sequence of the reads mapped to the European sardine complete mitochondrial genome (Genbank accession number GCF_963854185.1) by using the option -doFasta 2 and removing positions with sequencing depth below 3X (-setMinDepth 3), with no sign of contamination in ANGSD (Korneliussen et al., 2014).

Beagle files with the nuclear genome positions of single nucleotide polymorphisms (SNPs) were produced by ANGSD (Korneliussen *et al*. 2014) using the following options: angsd - bam $bamList -ref $REF -out ${out}.snp -C 50 -baq 2 -remove_bads 1 -uniqueOnly 1 - doCounts 1 -doGlf 2 -GL 1 -doMaf 2 -SNP_pval 1e-6 -doMajorMinor 1 -minQ 20 - minMapQ 30 -minInd 62 -minMaf 0.05 -rmTrans 1. Transitions were excluded. A restrictive minimum allele frequency of 0,05 was used to discard possible contamination. A total of 54,438 SNPs were obtained using all samples allowing for 50% missing data (-minInd62) and keeping only the SNPs that had information for at least one of the five samples with the lowest depth (1004, 1009, 1013, 1014, 1018, the ones that were not chosen for deep sequencing due to lower endogenous values). Admixture proportions were estimated by running NGSadmix version 32 (Skotte *et al*. 2013) for K equal 2 and 3 with 300 seed values, ensuring convergence. A principal component analysis (PCA) using the same SNP set was obtained with PCAngsd version 0.1 (Meisner & Albrechtsen 2018).

### (e) Authentication and alignment

To authenticate the historical sequences obtained as genuine, we ran DamageProfiler v0.4.9 (Neukamm *et al*. 2021) in order to detect DNA post-mortem damage patterns typical of ancient or degraded DNA. The program uses misincorporation patterns, particularly deamination of cytosine into uracils, within a Bayesian framework. An elevated C to T substitution rate towards sequencing starts (and complementary G to A rate towards the end) is considered indicative of genuine ancient or degraded DNA.

### (f) Species ID and Phylogenetic Analysis

Full mitochondrial DNA genomes were first aligned with mitogenomes from the European sardine, Atlantic chub mackerel (*Scomber colias*), Atlantic horse mackerel (*Trachurus trachurus*), anchovies (*Engraulis encrasicolus*), sprat (*Sprattus sprattus*), European seabass (*Dicentrarchus labrax*) and gilthead seabream (*Sparus aurata*).

Protein coding genes (except for ND6) and ribosomal RNA genes, as annotated in the European sardine reference mitochondrial genome, were extracted from the alignment and served as input in Beast 2.7.6 (https://doi.org/10.1371/journal.pcbi.1006650). Genes were input separately under a linked tree and clock model, and independent sites models, allowing each gene to have independent mutation rates. We picked a GTR nucleotide substitution model with a proportion of invariable sites, and a strict clock model, as we were more interested in taxonomic classification of our samples than splitting times. We picked the most complex model (GTR), as recommended by Huelsenbeck and Rannala (2004), as Bayesian phylogenetics is less sensitive to overspecification than the reverse. The MCMC chain ran from 10 million generations and 10% were discarded as burn in.

Historical mitogenomes from Adro Vello were aligned with 108 modern specimens from across the species distribution range (Barry *et al*. 2022; da Fonseca *et al*. 2024). PopART (Leigh & Bryant 2015) was used to estimate a median-joining haplotype network (Bandelt *et al*. 1999) using mitochondrial SNPs with minor allele frequency >25% (total of 30 SNPs), we adopted this conservative filter to decrease noise in the obtained network.

## Results and Discussion

### (a) Radiocarbon dating

The sample showed good preservation indicated by a C:N ratio of 3,3 which falls within the expected range (2,9-3,6) for well-preserved collagen (DeNiro 1985). The radiocarbon dating yielded an age of 2280±30 BP (Beta – 663761). The stable isotope values for carbon and nitrogen, ∂13C (-12,2) and ∂15N (8,3) align with expectations for a marine pelagic fish species (Fuller *et al*. 2012). After calibration this corresponds to a date range of 84-394 AD with 95,4% probability and 162-321 calibrated with 68,2% probability. The marine reservoir effect is a problem when dating marine organisms, it reflects the complex interaction between the atmosphere and the ocean in the global carbon cycle. This correction is especially important when comparing marine and terrestrial samples. Oceans are vast carbon reservoirs but because of complexities in marine circulation the actual reservoir effect correction varies with location. Northwestern Iberia is an active upwelling area, but the upwelling intensity varies depending on the latitude, and it may have changed throughout the Holocene. This regional difference from the average global marine reservoir correction is designated ΔR (Stuiver & Braziunas 1993). There are no ΔR values for Adro Vello, Galicia (lat 42.4775873, long -8.9313708) so we used the data from the 10 nearest points we found at http://calib.org/marine/ (Cook *et al*. 2015), which are all situated in the Portuguese Atlantic coast and we got a ΔR value of 12±163. The uncertainty of this value is however massive when compared to the correction factor. Applied to our radiocarbon date using the correction factor only we would have a range102-409AD if the uncertainty is taken into acount we get and age range of 161BC to 637AD. However, we also have information from the startigraphic study of the site which tells us the construction of the fish-salting plant can be dated to the beginning of the 1st century AD due to the finding of fragments of Haltern 70 amphorae in its walls. Haltern 70 is a type of Roman clay amphora of Baetican production, characteristic of the first half of the 1st century in Northwestern Iberia . In addition, a deposit excavated in 2023 directly linked to the factory has provided us with archaeological materials dating from the 1st, 2nd and early 3rd centuries AD (compositional date). Therefore, the abandonment and collapse of the factory must have taken place sometime in the early 3rd century AD. A similar date has been attributed to the salting factory located in O Naso beach (illa de Arousa, Pontevedra) (Fernández Fernández *et al*. 2022).

We must therefore attribute to the fish remains discovered in the interior of Vat 1 a date around the beginning of the 3rd century AD, in accordance with the 14C dating.

### (b) Extraction and library construction

All samples (18 bones), both crushed or simply submerged in the digestion buffer, were digested and yielded usable DNA, and we were able to build libraries for all of them. Of these, 17 specimens were sequenced and all yielded sardine DNA with a percentage of endogenous content between 4 and 21%, less than what has been described in the literature for bigger fish like the Atlantic cod (Ferrari *et al*. 2021). However, taking into account the special processing these samples endured, where whole fat fish were crushed and fermented in brine for a substantial period of time to produce *garum* or liquamen, allowing it to undergo enzymatic and bacterial action these are outstanding figures. The pH of these solutions was likely highly acidic, due to the fermentation process, which contributes to the acceleration of DNA degradation through time resulting in lower levels of endogenous DNA and a higher fragmentation rate. The combination of low pH and organic acids (especially lactic acid) is the main preservation factor in fermented fish products. Generally, pH should be below 5–4.5 in order to inhibit pathogenic and spoilage bacteria. Our findings, reveal new applications for these type of samples in the study of historical economies and past societies.

### (b) Species Identification

Twelve of the eighteen samples were sequenced to a higher depth and mapped against the European sardine genome, as they were all previously morphologically identified as sardines. After removing duplicates and presumed paralogs we were able to assemble their full mitogenome (17,552bp) with a mean depth of coverage between 1.3 and 197.6X (Table 1) (SRA project XXX). The nuclear genomes have depths of coverage between 1.1x and 3.7x. The damage patterns, as visualized using DamageProfiler v0.4.9, show an increase in the frequency of thymine and adenine in the 5’ and the 3’ end, respectively, confirming the endogenous nature of the DNA (Supplementary figure 1).

**Table 1.** – Sample information and mapping statistics for the 17 samples screened.

| Sample id | Bone | Sample weight (g) | % endogenous DNA | Reads after filtering | Reads mapped mtDNA | Average depth of coverage mtDNA | Reads mapped to the nuclear genome | Average genome depth of coverage | Average read length |
| --- | --- | --- | --- | --- | --- | --- | --- | --- | --- |
| 1001 | 2 articulated vertebrae | 0.021 | 16 | 141 710 149 | 29 880 | 119.4 | 16 613 665 | 2,4 | 74 |
| 1002 | 3 articulated vertebrae | 0.025 | 15 | 151 183 746 | 12 444 | 48.1 | 8 063 237 | 1,9 | 80 |
| 1003 | vertebrae | 0.010 | 20 | 168 636 510 | 24 869 | 95.1 | 19 663 304 | 2,6 | 71 |
| 1004 | vertebrae | 0.010 | 4 | 2 175 980 | 301 | 1.8 | 79 495 | 1,1 | 70 |
| 1005 | vertebrae | 0.003 | 25 | 31 453 540 | 10 474 | 38.7 | 6 777 841 | 1,6 | 71 |
| 1006 | vertebrae | 0.009 | 23 | 177 387 467 | 46 293 | 197.6 | 29 736 874 | 3,7 | 72 |
| 1007 | 2 articulated vertebrae | 0.018 | 0 | 0 | 0 | 0 | 0 | 0 |  |
| 1008 | vertebrae | 0,022 | 15 | 140 235 210 | 17 154 | 67.2 | 17 111 757 | 2,5 | 78 |
| 1009 | vertebrae | 0.020 | 9 | 1 714 207 | 211 | 1.6 | 132 402 | 1,1 | 78 |
| 1010 | vertebrae | 0.015 | 14 | 169 175 081 | 13 999 | 57.1 | 12 777 748 | 2,3 | 83 |
| 1011 | vertebrae | 0.016 | 15 | 163 644 799 | 4 291 | 17.8 | 3 732 925 | 2,1 | 87 |
| 1012 | vertebrae | 0.022 | 17 | 182 394 811 | 12 535 | 52.6 | 10 961 754 | 2,3 | 84 |
| 1013 | vertebrae | 0.032 | 8 | 1 715 636 | 181 | 1.6 | 126 096 | 1,1 | 78 |
| 1014 | vertebrae | 0.022 | 9 | 2 255 511 | 317 | 1.8 | 182 473 | 1,1 | 78 |
| 1015 | 2 articulated vertebrae | 0.020 | 15 | 114 584 604 | 17 414 | 70.5 | 7 991 159 | 1,9 | 82 |
| 1016 | vertebrae | 0.010 | 23 | 141 614 267 | 30 479 | 128.3 | 21 489 609 | 3,0 | 77 |
| 1017 | operculum | 0.033 | 13 | 160 855 814 | 21 496 | 86.8 | 15 959 266 | 2,4 | 80 |
| 1018 | operculum | 0.043 | 6 | 1 760 720 | 105 | 1.3 | 87 091 | 1,1 | 68 |

Species identification was also confirmed by the Bayesian phylogenetic reconstruction (Figure 2A). The 12 specimens from Adro Vello form a clade with the reference mitogenome of the European Sardine with a posterior probability of one (pp=1). The median joining network constructed using 120 individuals from across the sardine distribution range show that our historical specimens mainly cluster with individuals from the Western and Central groups (da Fonseca *et al*. 2024), showing haplotype continuity in the last 2000 years in this region.

### (c) Sardine population structure

We compared the nuclear data obtained from Adro Vello specimens to the genomes of modern individuals collected from across the species distribution range previously published by da Fonseca et al (2024) and Barry et al (2022).

The admixture analyses conducted in NGSadmix showed that our Adro Vello population clusters with the previously described Centre cluster. This group encompasses samples from Morocco, Iberia and the Atlantic coast of France. Notoriously these samples seem much less admixed than their modern counterparts from Western Iberia, with only a sample presenting a small contribution from the eastern group.

The principal component analysis also places our samples in the Centre cluster with the first component explaining 7.3% of the variation and PC2 1,2%. The samples from Adro Vello form a group. The lower admixture rates and greater differentiation from both Mediterranean and the Macaronesian populations suggest reduced connectivity for this species in Roman times.

The current scenario, characterized by increased admixture rates in modern populations, may result from human activities such as intensive fishing and shipping movement in the area. The European sardine is a very important economic fish species, especially in Southern Europe and Morocco, being the main target of the purse-seine fleets in Portugal and Spain. This species represents a major source of income for local economies and has been overexploited in the last decades (ICES 2013).

### (d) Resource use in O Grove (Galicia) and the Southern Atlantic façade

The European sardine was the main fish resource used for the making of fish sauce in the archaeological site of O Grove, Galicia and in the other fish salting plants identified on the coast of Galicia (González Gómez de Agüero 2014), but also in those located on the rest of the Atlantic façade.

For example, on the French Atlantic coast, in a roman fish-salting plant identified on Rue du Guet (Douarnenez, France), production residue based on sardine (*Sardina pilchardus*) and some sparids (gilthead seabream) was documented (Desse & Desse 1983). In Brittany (NW France), a residue of *Sardina pilchardus* was identified in the Douarnenez II fish-salting plant (Sanquer 1977) as well as in the Lanévry plant (Kerlaz and Driard, 2011) and in La Falaise site (Étel). In the latter, in vat number one, 99,6% of the halieutic remains were *Sardina pilchardus* and the rest (0,4%) were a mix of Atlantic herring (*Clupea harengus*), whiting (*Merlangius merlangus*) and Atlantic mackerel (*Scomber scombrus*) (Ephrem, 2016). On the coast of Portugal, we have more examples of the preponderance of sardine in Roman industrial contexts. In Lisbon, on Rua Augusta, a fish-salting factory was excavated and the ichthyological residue recovered inside the vats had a clear prevalence of *Sardina pilchardus* (Fabião 2017). Nearby, in Belém, a large fish-salting plant was excavated in the old Governor’s House. Remains of ichthyofauna were recovered from 16 vats and *Sardina pilchardus* represented more than 98% of the recovered material with other residual species such as anchovies (*Engraulis encrasicolus*), Atlantic horse mackerel (*Trachurus trachurus*), Atlantic mackerel (*Scomber scombrus*) and Mediterranean moray (*Muraena helena*) also present (Fabião *et al*. 2021). Further south, in the large fish producing centre of Tróia (Setúbal) the presence of *Sardina pilchardus* was documented in Tróia 1 factory along with clupeids, *Acipenseridae*, axillary seabream (*Pagellus acarne*) and Atlantic mackerel (Étienne *et al*. 1994) .

In Baelo Claudia, a Roman city in Baetica, close to the Strait of Gibraltar, several kinds of *garum* were prepared. In some *cetariae*, *Engraulis encrasicolus* was the main species used while in others 95% of the remains were from European sardine. Besides sprat and sardine *garum*, axillary seabream *garum* was also produced (Bernal-Casasola *et al*. 2018). In the Atlantic territory of North Africa, the situation seems similar. In Tahaddart (Assilah, Morocco) sauces based on sardine and a clupeid, most abundant resource, have been identified (Adolfo Fernández, personal communication).

## Conclusions

A great advantage of using samples from the bottom of fish salting vats it is their contemporaneity ensuring us to be sampling the same population, often a huge problem in ancient DNA studies.

In cases where bones are extremely crushed and therefore morphologically unidentifiable this methodology or bulk bone metabarcoding can be used as a complimentary method for fish species identification in a similar fashion to what has been previously done for terrestrial animals (Grealy et al., 2015; Murray et al., 2013).

In this study, we demonstrate that usable DNA can survive in fermentation environments, such as the brines used by the Romans to make *garum*, allowing for species identification and the study of ancient populations. To our knowledge this is the first time DNA has been extracted and sequenced from remains from the bottom of Roman fish salting vats, allowing for species identification of crushed bones, opening a new window into the past for the study of archaeological fish remains, a material that has been often overlooked. This approach would work for many other fish species that have a reference genome/ mitogenome, paving the road to systematic studies of fish species used in different sites, eras or cultures and its population dynamics.

## Acknowledgements

This research was partially supported by the Strategic Funding UIDB/04423/2020 and UIDP/04423/2020, through national funds provided by FCT and European Regional Development Fund (ERDF), in the framework of the programme PT2020. PFC was partially supported by national funds through FCT, I.P., under the Scientific Employment Stimulus Initiative, references CEECIND/01799/2017 and 2023.05877.CEECIND, SARDINOMICS 2022.03142.PTDC and the European programme SYNTHESYS+ (SE-TAF-TA3-005), which funded the visit and access to a clean lab facility at NHM Stockholm, University of Stockholm. We would also like to thank Professor Love Dalén for hosting PFC in his ancient lab and Edana Lord and Vendela Kempe Lagerholm for lab guidance.

---

Supplementary Figure 1-Map damage profiles for samples 1001-1018 (except 1007).

