## Supplementary figures and images for "DNA from *cetariae* fish remains confirms sardine (*Sardina pilchardus*) use and local population continuity in Northwestern Iberia since Roman times"

### Supplementary Figure 1

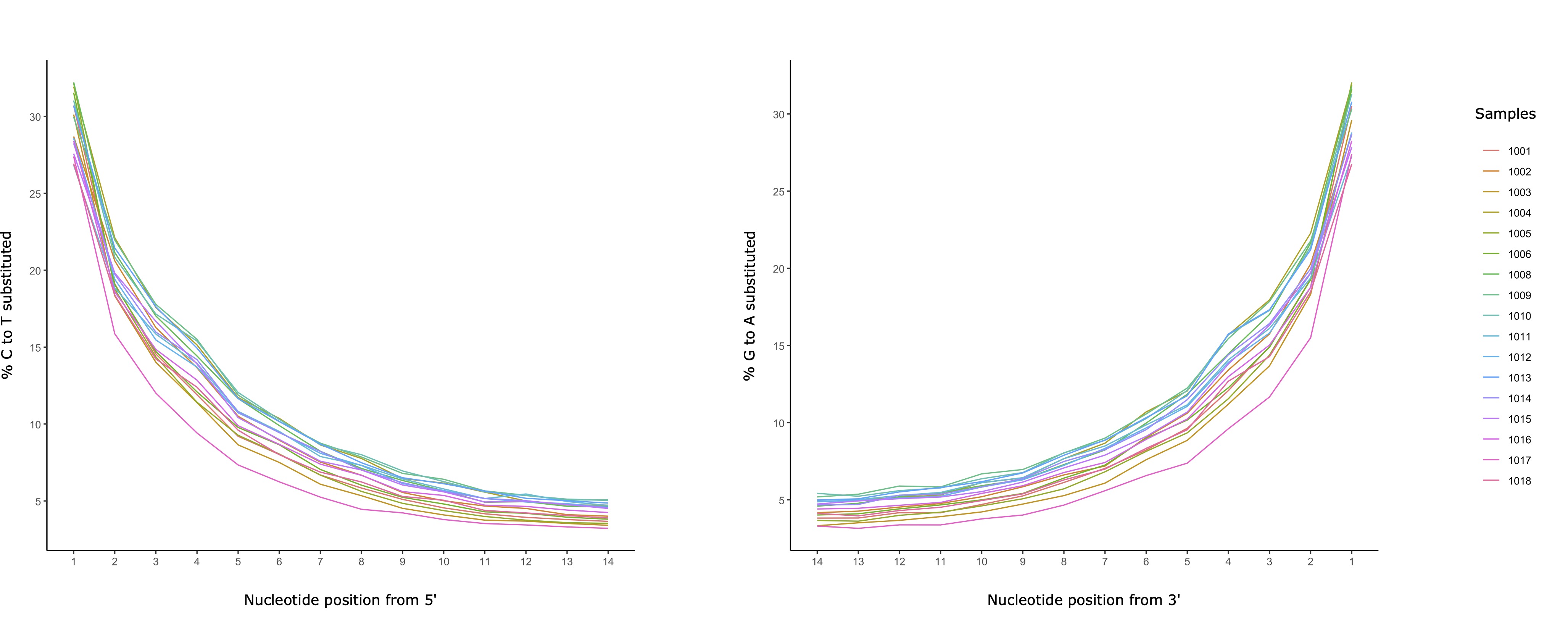
